# Highly Efficient Repair of the ΔF508 Mutation in Airway Stem Cells of Cystic Fibrosis Patients with Functional Rescue of the Differentiated Epithelia

**DOI:** 10.1101/561183

**Authors:** Sriram Vaidyanathan, Ameen A. Salahudeen, Zachary M. Sellers, Dawn T. Bravo, Shannon S. Choi, Arpit Batish, Wei Le, Sean De La O, Milan P. Kaushik, Noah Galper, Ciaran M. Lee, Gang Bao, Eugene H. Chang, Jeffrey J. Wine, Carlos E. Milla, Tushar J. Desai, Jayakar V. Nayak, Calvin J. Kuo, Matthew H. Porteus

**Affiliations:** Department of Pediatrics, Stanford University, Stanford, CA 94304, USA; Department of Hematology, Stanford University, Stanford, CA 94305, USA; Department of Otolaryngology-Head and Neck Surgery, Stanford, CA 94305, USA; Department of Bioengineering, Rice University, Houston, TX 77030, USA; Department of Otolaryngology, University of Arizona – Tucson, Tucson, AZ 85724, USA; Department of Psychology, Stanford University, Stanford, CA 94305, USA; Department of Pulmonary and Critical Care Medicine, Stanford University, Stanford, CA 94305, USA

## Abstract

Cystic fibrosis (CF) is a monogenic autosomal recessive disorder caused by mutations in the Cystic Fibrosis Transmembrane Conductance Regulator (CFTR) Cl^-^ channel. CF results in multiorgan dysfunction and ultimately mortality from respiratory sequelae. Although pharmacologic approaches have demonstrated efficacy in reducing symptoms and respiratory decline, a curative treatment modality remains elusive. Gene therapy, a promising curative strategy, has been limited due to poor correction efficiencies both *in vitro* and *in vivo*. Here, we use Cas9 and adeno-associated virus 6 (AAV6) to correct the ΔF508 mutation (found in ∼70% of CF alleles and ∼90% of CF patients in North America) in upper airway basal stem cells (UABCs) obtained from CF and non-CF patients undergoing functional endoscopic sinus surgery (FESS). In UABCs from homozygous (ΔF508/ΔF508) and compound heterozygous (ΔF508/Other) CF patients, we achieved 28 ± 5 % and 42 ± 15% correction, respectively. In homozygous human bronchial epithelial cells (HBECs), we achieved 41± 4 % correction. Upon differentiation in air-liquid interface (ALI), cultures of corrected CF cells displayed partial restoration of CFTR_inh_-172 sensitive Cl^-^ currents relative to non-CF controls: 31± 5 % in UABCs and 51 ± 3 % in HBECs (both from subjects homozygous for ΔF508 CFTR). Finally, gene edited cells embedded successfully and retained expression of cytokeratin 5 (KRT5), a basal cell marker, on a FDA-approved porcine small intestinal submucosal (pSIS) membrane previously shown to improve re-mucosalization after FESS. In summary, we present an efficient, feeder-free, selection-free and clinically compatible approach to generate cell-based therapies for CF from autologous airway stem cells. This approach represents a first step towards developing patient-specific autologous airway stem cell transplant as a curative treatment for CF.

## Introduction

Cystic fibrosis (CF) is an autosomal recessive monogenic disease caused by mutations in the Cystic Fibrosis Transmembrane Conductance Regulator (CFTR) protein, a Cl^-^ channel. Although CF is a systemic disease that affects exocrine function in multiple organ systems, CF lung disease is the major cause of morbidity and mortality in CF patients. Over the past decade, small molecule CFTR correctors and potentiators have been developed, and represent a significant advancement in CF therapeutics (Van Goor et al., 2009, 2011). Although these small molecules improve lung function and reduce pulmonary exacerbations in patients, they are expensive, show variable therapeutic responses, have significant adverse effects (e.g. hepatotoxicity, dyspnea), must be administered daily (Wainwright et al., 2015) and necessitate lifelong treatment. As a result of these limitations, there is continued interest in developing genome editing strategies to correct CFTR mutations in stem cells for achieving durable restoration of native CFTR function.

Genome editing using zinc-finger nucleases or CRISPR/Cas9 has been attempted in intestinal stem cells and induced pluripotent stem cells (iPSCs), respectively (Crane et al., 2015; Firth et al., 2015; Schwank et al., 2013). Genome editing using nucleases involves the creation of a double-stranded break (DSB) which is then repaired using either the non-homologous end joining (NHEJ) or homologous recombination (HR) pathways. NHEJ results in insertions and deletions (INDELs) and cannot be used to precisely correct mutations. The HR pathway can be used to correct mutations but it has been challenging to achieve high levels of HR to correct CFTR mutations in primary stem cells. Previous studies focused on the ΔF508 mutation that affects >70% of CFTR patients and employed selectable markers to enrich for cells corrected using the HR pathway. The efficiencies reported in these studies (0.02% (Schwank et al., 2013) before selection and 16% after selection (Crane et al., 2015)) are useful to understand the pathophysiology of different mutations and may enable drug screening, but are too low for clinical utility. In addition to low correction frequencies, subsequent directed differentiation of iPSCs to produce a clinically relevant airway cell type remains a major challenge and the risk of teratoma formation from residual undifferentiated iPSCs in the population remains. In contrast, a highly efficient, selection-free, feeder-free, genome editing strategy in endogenous airway stem cells could facilitate the rapid development of durable cell therapies to treat CF airway disease.

We describe experiments using Cas9 and adeno-associated virus 6 (AAV6) to correct the ΔF508 mutation using the HR pathway in *ex-vivo* expanded, human upper and lower airway cytokeratin 5+ (KRT5+) stem cells from the upper airway and bronchial epithelium, respectively. We pursued this *ex-vivo* correction strategy to overcome several obstacles associated with *in vivo* gene correction including successful vector delivery across the thick airway mucus barrier (Armstrong et al., 2014), and the immunogenicity to Cas9 that is highly prevalent in humans (Charlesworth et al., 2019; Wagner et al., 2018) and develops in animals after exposure to Cas9 *in vivo* (Wang et al., 2015a).

Recent studies have achieved >50% allelic modification in hematopoietic stem cells, T-cells, and embryonic stem cells using Cas9 with single guide RNA (sgRNA) modified with 2’-O-methyl 3’phosphorothioate (MS) in the 5’ and 3’ terminal nucleotides (MS-sgRNA) and correction templates delivered by AAV6 (Bak et al., 2018; Dever et al., 2016; Hendel et al., 2015; Martin et al., 2018). We modified and adapted this general strategy to airway stem cells. We present the following findings:

1. Correction of the ΔF508 mutation in 28 ± 5% and 42 ± 15% alleles in KRT5+ basal stem cells derived from the upper airway epithelium of ΔF508 homozygous and compound heterozygous patients, respectively. In addition, we achieved 41 ± 4% correction in KRT5+ stem cells obtained from bronchial epithelia (HBECs) of ΔF508 homozygous patients.
2. Corrected KRT5+ upper airway basal stem cells (UABCs) from homozygous CF patients differentiated in air-liquid interface showed 31 ± 5% CFTR_inh_-172 sensitive Cl^-^ short-circuit currents relative to non-CF cultures subject to the same assay (Ma et al., 2002; Vu et al., 2017). By contrast, corrected KRT5+ HBECs showed 51 ± 3% CFTR_inh_-172 sensitive Cl^-^ short-circuit currents relative to non-CF HBEC cultures subjected to the same assay.
3. Gene edited KRT5+ UABCs were successfully embedded on a FDA-approved porcine small intestinal submucosal (pSIS) surgical matrix routinely used for epithelial repair during functional endoscopic sinus and skull base surgeries (FESS) (Nayak et al., 2018).

In addition to providing a highly efficient, clinically compatible approach for correcting the ΔF508 mutation, our experiments indicate that even mosaic correction of 50% or less of the stem cell population can achieve significant restoration of CFTR function within the *in vitro* differentiated epithelium. Combined with successfully embedding corrected cells on a FDA-compatible scaffold that is already in use for sinonasal repair, these experiments form the foundation for a strategy of transplanting genetically corrected autologous airway stem cells embedded in a matrix back into the airways of patients to potentially cure the respiratory complications of CF.

## Results

### Isolation and culture of UABCs

Cas9-based gene editing utilizing the homologous recombination pathway to correct mutations has been shown to be improved under conditions that promote cell proliferation (Charlesworth et al., 2018). Therefore, we first optimized cell culture conditions to both increase proliferation of basal cells and improve their survival after electroporation (also called nucleofection).

We obtained sinonasal tissue from non-CF and CF patients undergoing FESS and performed pronase digestion and red blood cell lysis. On day zero, 2-22% (mean ± S.D. = 10 ± 8%) cells were found to express KRT5+, a marker for stem/progenitor cells in upper and lower airway epithelia (Figure S1A) (Bravo et al., 2013; Wang et al., 2015b). We initially cultured UABCs as 3-dimensional organoids without any feeder cells in Matrigel domes. Epidermal growth factor (EGF) along with the bone morphogenetic protein (BMP) antagonist Noggin were sufficient to promote organoid growth, alongside, the transforming growth factor-β (TGF-β) inhibitor A83-01, and the Rho-kinase inhibitor Y-27632. Such UABC organoids submerged under media were KRT5+ (Fig. S1B-E). Under these feeder-free organoid conditions, KRT5+ UABCs were enriched after 5 days in culture from 10 ± 8% to 72 ± 18 % (Figure S1A). Optimal cell density was empirically determined to be ∼10,000 cells/cm^2^ (Figure S1F).

Notably, human UABC could also be cultured and expanded as KRT5+ monolayers using the same EGF/Noggin media (EN media) used for culturing organoids (Fig. S2A). Using cells from the same donors as in Figure S1F, optimal cell density was empirically determined to be ∼10,000 cells/cm^2^ for cells cultured as monolayer and resulted in a similar fold-expansion of KRT5+ cells (Figure S2A). We attempted gene editing on UABCs cultured as organoids and cells cultured as monolayers using electroporation of a previously reported control homologous recombination (HR) template (or donor template) expressing green fluorescent protein (GFP) at the CCR5 locus (Hendel et al., 2015). Since HR was more efficient in basal cells cultured as monolayers (Fig. S2B), we used this approach in subsequent experiments with the EN media optimized from organoids. Optimal cell density was also tested for cells passaged as monolayers and was empirically determined to be ∼10,000 cells/cm^2^ at passage 1. (Figure S2C). Cell culture at 5% O_2_ also improved cell proliferation of KRT5+ UABC monolayers compared to 21% O_2_ (Figure S2D).

### Insertion of Correction Sequence by Homologous Recombination in ΔF508 Locus

Several sgRNAs in the exon carrying the ΔF508 mutation (exon 11) were tested and the most efficient sgRNA (Figure 1A) was used for further experiments. This sgRNA is active even in non-ΔF508 cells since the protospacer sequence is eight base pairs away from the ΔF508 mutation. AAV6 at a multiplicity of infection (MOI) of 10^6^ particles per cell was found to have the highest transduction among commonly used serotypes (Figure S3A-B). Silent mutations were introduced in the correction templates to both prevent re-cutting by Cas9 and to serve as a marker to quantify insertion in non-ΔF508 cells. We tested the efficiency of two correction sequences carrying different silent mutations on HR (Figure S3C). In these experiments, the CFTR exon 11 locus was amplified using junction polymerase chain reaction (PCR) 4 days after editing, after which insertions and deletions (INDELs) and HR events were quantified using TIDER (Brinkman et al., 2018). We found that a correction sequence with 6 silent mutations surrounding the Cas9 double-stranded break was more effective than one with 4 silent mutations (Figure S3C). Further experiments were thus performed with the codon diverged sequence carrying 6 silent mutations (Figure 1A).

**Figure 1:**
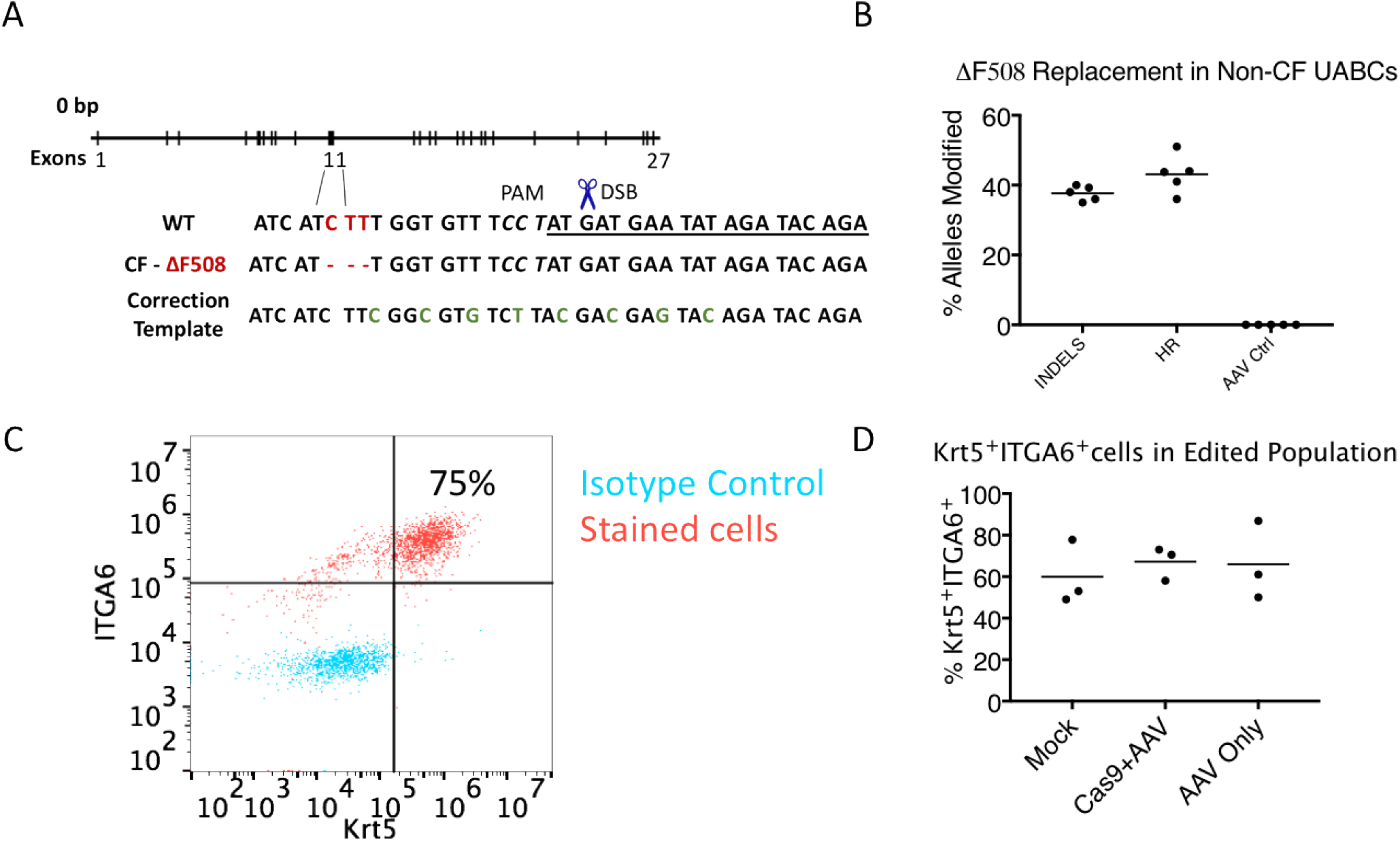
**(A)** Schematic describing the Cas9/AAV mediated strategy to correct ΔF508. The underlined segment represents the sequence complementary to sgRNA used. The PAM (protospacer adjacent motif) is indicated in italics for the wild-type (WT) sequence. Silent mutations introduced in the correction template to prevent re-cutting by Cas9 are colored in green. **(B)** The region around exon 11 was amplified using IN-OUT PCR to quantify INDELS and HR using TIDER. INDELs were observed in 38 ± 2 % alleles and HR was observed in 43 ± 5% alleles. Controls treated with only AAV did not show any INDELs or HR. **(C)** On day 4 after editing, the UABCs were stained for KRT5 and Integrin alpha 6 (ITGA6). A representative FACS plot shows that 75% of edited cells were KRT5+ITGA6+^·^**(D)** In trials on three different donors, the KRT5+ITGA6+ population was similar between control and edited UABCs.

Using the optimal template, our correction sequence was observed in 43 ± 5% alleles (Figure 1B) and INDELs were observed in 38 ± 2% of the alleles (Figure 1B), a ratio of HR:INDEL of >1 which is consistent with prior work that when optimal amounts of donor are delivered to dividing cells, high frequencies of HR-mediated editing are possible (Charlesworth et al., 2018; Hendel et al., 2014). On the day of extraction, 67 ± 8% of edited cells were KRT5+ and Integrin Alpha-6+ (ITGA6+) which was similar to control (mock) cells, with 60 ± 15% (Figure 1C-D). Thus, insertion of the correction sequence did not alter the phenotype of the targeted basal cells (Bravo et al., 2013; Wang et al., 2015b).

### Off-target activity

Cas9 nucleases can tolerate mismatches between the sgRNA and genomic DNA sequences. This results in unintended off-target DSB depending on the number, position and distribution of mismatches (Hsu et al., 2013; Lin et al., 2014). Such off-target activity is undesirable as it can result in mutations in important genes (e.g., oncogenes). *In-silico* methods predicted over 50 possible off-target sites. The off-target activity of the MS-sgRNA was characterized at the top 47 predicted off-target sites (Table S1). Off-target activity above background levels (0.1%) was observed in only one site. INDELs were observed in 0.17% of alleles in Chr11:111971753-111971775 (OT-41 in Table S1). This region corresponds to an intron of the gene coding for the protein DIXDC1. DIXDC1 is a regulator of Wnt signaling and has been shown to be active in cardiac and neural tissue but not in airway cells (Wang et al., 2006). The intronic target sequence is of no known functional significance and does not occur in the putative splice donor or acceptor sequences of the intron.

### Correction of ΔF508 mutation in KRT5+ stem cells expanded from primary CF patient airway epithelia

We used the optimized protocol to correct the ΔF508 mutation in both UABCs and HBECs from homozygous (ΔF/ΔF) patients and UABCs from compound heterozygous (ΔF/other) patients. In UABCs and HBECs from homozygous patients, we observed allelic correction rates of 28 ± 5% and 41± 4% alleles, respectively. We observed 42 ± 15% allelic correction in compound heterozygous UABC samples (Figure 2A). Gene-corrected airway cells cultured in air-liquid interface (ALI) using Transwells retained their ability to generate a pseudostratified epithelium with a basal layer of KRT5+ cells and a luminal layer of ciliated cells (acetylated α-tubulin+) and mucin secreting cells (Muc5AC+) (Figure 2B-C). The fraction of edited cells did not change appreciably over the 28-35 days during which cells were cultured in ALI, suggesting equal contribution of corrected and uncorrected basal cells to reconstituting the epithelium (Figure 2D). Figure 2E shows a representative Western blot probing CFTR expression in non-CF, uncorrected and corrected CF samples after differentiation in ALI (CFTR Antibody 450). Mature CFTR expression was not observed in the uncorrected homozygous sample (lane 3) and was restored in cells corrected using the Cas9/AAV6 platform (lane 4). Mature CFTR expression in corrected cells was appreciable, although less than that seen in non-CF nasal cells (lane 2). These results indicate that gene edited basal cells can reconstitute an epithelium with restored CFTR protein expression *in vitro*.

**Figure 2:**
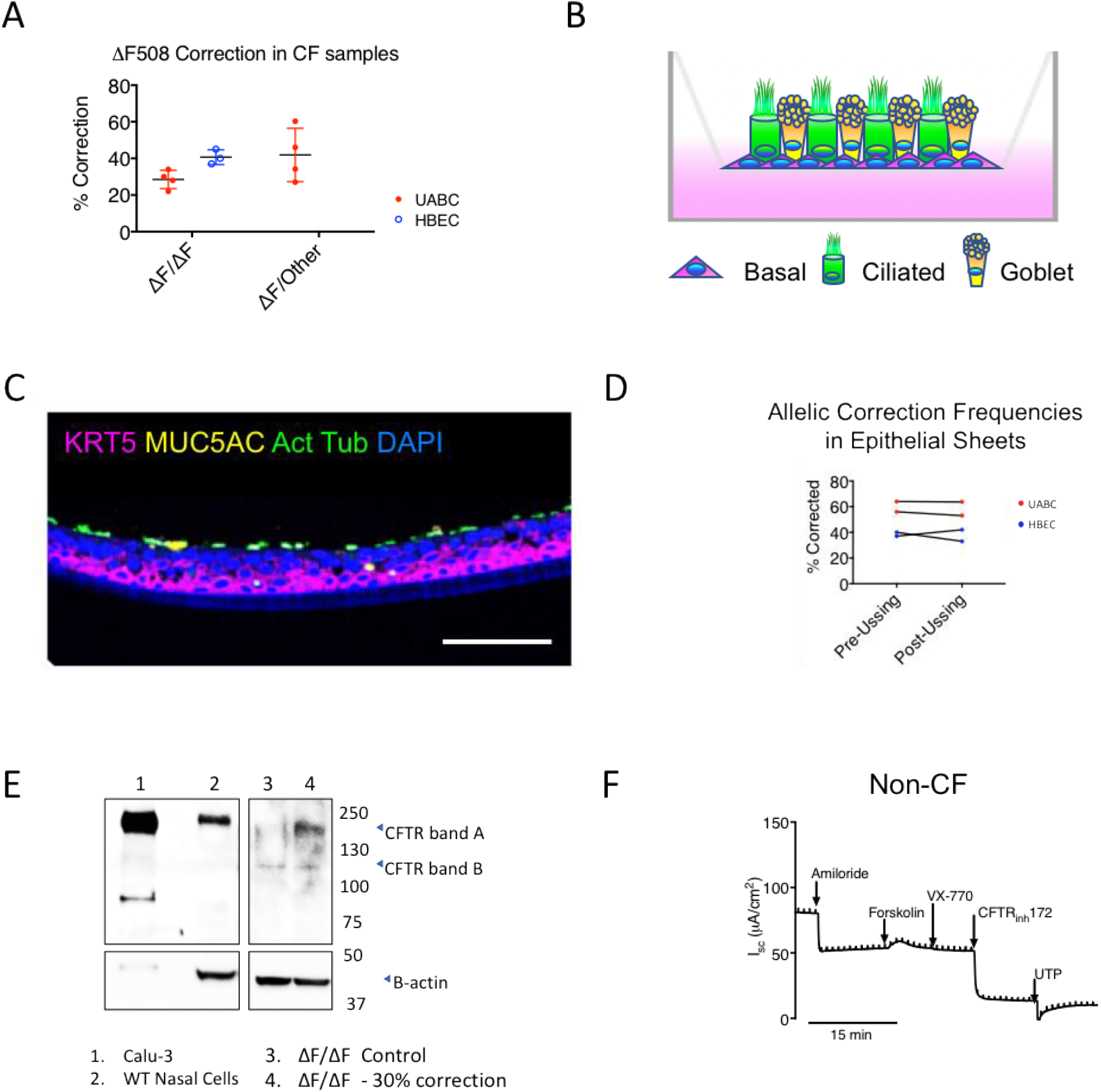
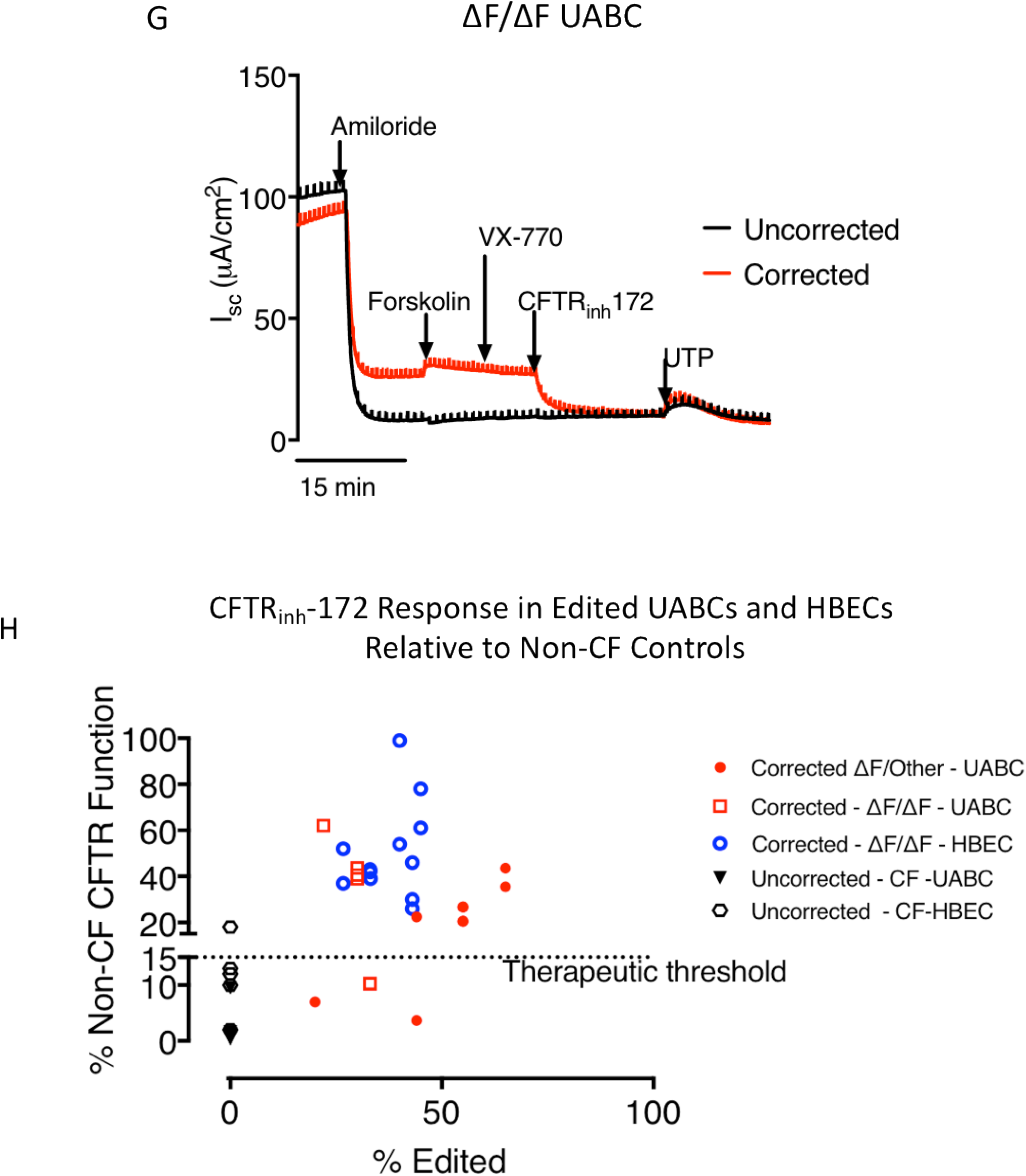
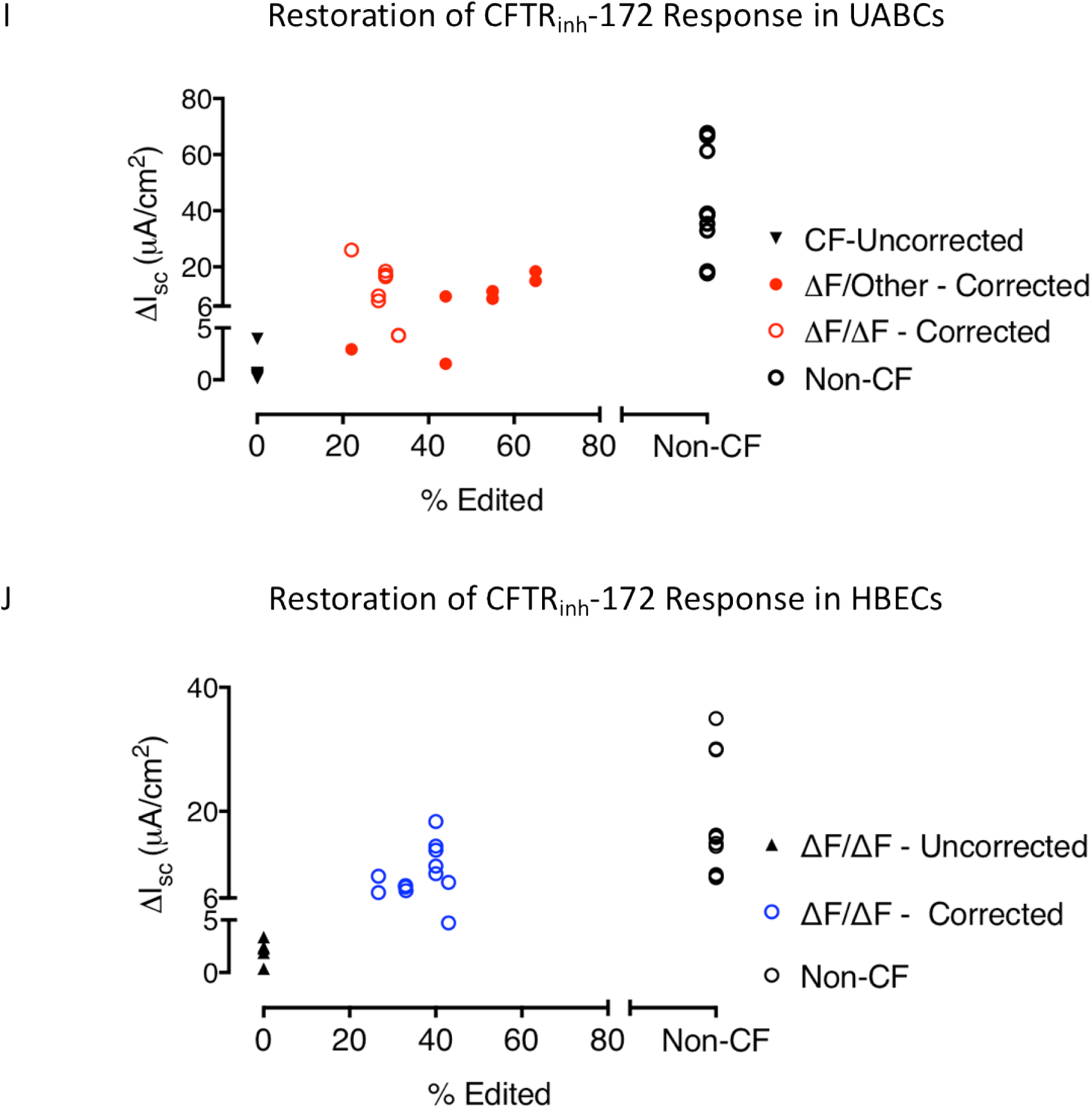
**(A)** Summary of percent alleles exhibiting HR in CF patient samples (ΔF/ΔF – homozygous and ΔF/other – Compound heterozygous) **(B)** Illustration of ideal ALI culture platform, with KRT5+ basal cells giving rise to overlying ciliated and goblet differentiated cell types **(C)** Edited CF cells cultured on ALI differentiate into a sheet with basal cells (KRT5+), ciliated cells (α-tubulin+) and mucus producing cells (Muc5AC+). **(D)** Allelic correction rates in edited cells were assessed at the time of plating on ALI and at the end of the Ussing assay 28-35 days later. Allelic correction rates as measured by TIDER did not change appreciably before and after culturing in ALI for Ussing assays, suggesting equal contribution of corrected and uncorrected basal cells to reconstituting the epithelium. **(E)** Western blot probing CFTR expression. Calu-3 cells were used as a positive control for mature CFTR (lane 1). Despite a lower amount of total protein being loaded in the gel, Calu-3 cells exhibited very high levels of CFTR expression. Calu-3 cells also showed lower expression of β-actin. Non-CF nasal cells (lane 2) showed a clear band corresponding to the higher molecular-weight mature CFTR (Band A). Mature CFTR expression was absent in non-edited ΔF508 homozygous (ΔF/ΔF) cells (lane 3) but a faint, lower molecular-weight band corresponding to immature CFTR was present (CFTR Band B). ΔF508 homozygous (ΔF/ΔF) cells after correction showed a restored mature CFTR band (lane 4) while also retaining a portion of the immature CFTR band. β-actin was used as a loading control. **(F)** Representative traces obtained from epithelial monolayers by Ussing chamber analysis in non-CF patients **(G)** Representative traces obtained from epithelial monolayers by Ussing chamber analysis in uncorrected and corrected ΔF508 homozygous samples. **(H)** CFTR_inh_-172 sensitive short-circuit currents observed in corrected CF samples as a percent of non-CF controls (UABCs: ΔF/ΔF: n = 4 donors, ΔF/other: n = 3 donors. HBECs: ΔF/ΔF: n = 3 donors). Studies in patients with milder mutations have suggested that even 15% CFTR function relative to non-CF subjects would be therapeutically beneficial (Char et al., 2014; Sheppard et al., 1993). **(I)** CFTR_inh_-172-sensitive short-circuit currents observed in non-CF, uncorrected and corrected CF UABC samples as a function of editing (ΔF/ΔF: n = 4 donors, ΔF/other: n = 3 donors and non-CF: n = 3 donors) **(J)** CFTR_inh_-172-sensitive short-circuit currents observed in non-CF, uncorrected and corrected CF HBECs as a function of editing (n = 3 donors, non-CF: n = 2 donors)

### Recovery of CFTR function in airway epithelial cells differentiated from gene-corrected basal cells

To evaluate CFTR function, we differentiated UABCs and HBECs (corrected CF, uncorrected CF and non-CF) in air-liquid interface cultures and measured currents in short-circuited monolayers using Ussing chamber assays. Representative traces from non-CF and CF epithelial monolayers from UABCs are shown in Figure 2F-G. Traces from epithelial monolayers derived from HBECs are shown in Figure S4. Consistent with expectations from Western blot results, corrected CF samples showed restored CFTR function, as indicated by increased CFTR_inh_-172-sensitive short-circuit current (Figure 2G). Note that the forskolin-stimulated currents were small for both control and corrected monolayers. Because culture conditions can activate CFTR to various extents, the magnitude of responses to inhibition by CFTR_inh_-172 provide a better indication of CFTR function than does the magnitude of the forskolin response. CFTR_inh_-172-sensitive short-circuit currents in corrected samples were lower than corresponding short-circuit currents in non-CF samples. The CFTR_inh_-172-sensitive currents in edited UABCs and HBECs are presented as a percent of non-CF controls in Figure 2H. Genotype information, percent alleles corrected and restoration of CFTR_inh_-172 short-circuit currents relative to average non-CF currents for individual samples are presented in Table 1. The CFTR_inh_-172-sensitive currents from corrected UABCs (n = 4 ΔF/ΔF donors; n = 3 ΔF/other; n = 3 non-CF) and HBECs (n = 3 ΔF/ΔF; n = 2 non-CF) are plotted as a function of allelic correction in Figures 2I and 2J, respectively. UABC cultures from compound heterozygotes with higher editing efficiencies showed correspondingly greater restoration of CFTR function. HBEC samples from 3 donors had similar correction rates and showed similar CFTR_inh_-172-sensitive currents.

**Table 1:** Summary of Percent Allelic Correction of (ΔF508) in CF UABC and HBEC Samples and Relative CFTR Function with Respect to Non-CF Controls

| Patient | Genotype | Percent Editing | Raw Inhibitable CF current (μA) | Percent Non-CF inhibitable Current | Cell Type | Sex | Age |
| --- | --- | --- | --- | --- | --- | --- | --- |
| **Non-CF Inhibitable Current UABC (Mean ± S.D.) = $42 \pm 6$** |  |  |  |  |  |  |  |
| Patient 1 | ΔF/Other | 22 | -2.93 | 7 | UABC | Male | 22 |
| Patient 2 | ΔF/Other | 44 | -1.55 | 5 | UABC | Female | 26 |
| Patient 2 | ΔF/Other | 44 | -9.5 | 22 | UABC | Female | 26 |
| Patient 3 | ΔF/Other | 66 | -18.41 | 45 | UABC | Male | 44 |
| Patient 3 | ΔF/Other | 66 | -15 | 35 | UABC | Male | 44 |
| Patient 3 | ΔF/Other | 54 | -8.67 | 21 | UABC | Male | 44 |
| Patient 3 | ΔF/Other | 54 | -11.27 | 27 | UABC | Male | 44 |
| Patient 3 | ΔF/Other | 54 | -11.32 | 27 | UABC | Male | 44 |
| Patient 3 | ΔF/Other | 54 | -8.71 | 21 | UABC | Male | 44 |
| Patient 4 | ΔF/ΔF | 30 | -17.16 | 41 | UABC | Male | 19 |
| Patient 4 | ΔF/ΔF | 30 | -16.46 | 39 | UABC | Male | 19 |
| Patient 4 | ΔF/ΔF | 30 | -18.45 | 44 | UABC | Male | 19 |
| Patient 5 | ΔF/ΔF | 33 | -4.27 | 10 | UABC | Unknown | Unknown |
| Patient 5 | ΔF/ΔF | 33 | -4.29 | 10 | UABC | Unknown | Unknown |
| Patient 6 | ΔF/ΔF | 22 | -26.04 | 76 | UABC | Male | 32 |
| Patient 7 | ΔF/ΔF | 28 | -9.7 | 28 | UABC | Unknown | Unknown |
| Patient 7 | ΔF/ΔF | 28 | -7.87 | 22 | UABC | Unknown | Unknown |
| **Non-CF Inhibitable Current HBEC (Mean ± S.D.) = $18 \pm 3$** |  |  |  |  |  |  |  |
| Patient 8 | ΔF/ΔF | 33 | -7.75 | 42 | HBEC | Male | 25 |
| Patient 8 | ΔF/ΔF | 26.7 | -6.87 | 37 | HBEC | Male | 25 |
| Patient 8 | ΔF/ΔF | 33.1 | -7.19 | 39 | HBEC | Male | 25 |
| Patient 8 | ΔF/ΔF | 43 | -5.51 | 30 | HBEC | Male | 25 |
| Patient 8 | ΔF/ΔF | 43 | -4.73 | 26 | HBEC | Male | 25 |
| Patient 8 | ΔF/ΔF | 43 | -8.52 | 46 | HBEC | Male | 25 |
| Patient 8 | ΔF/ΔF | 26.7 | -9.52 | 52 | HBEC | Male | 25 |
| Patient 8 | ΔF/ΔF | 33 | -8 | 43 | HBEC | Male | 25 |
| Patient 9 | ΔF/ΔF | 40 | -9.91 | 54 | HBEC | Male | 32 |
| Patient 9 | ΔF/ΔF | 40 | -18.36 | 99 | HBEC | Male | 32 |
| Patient 10 | ΔF/ΔF | 45 | -13.71 | 61 | HBEC | Male | 44 |
| Patient 10 | ΔF/ΔF | 45 | -11.17 | 78 | HBEC | Male | 44 |

Corrected ΔF508 homozygous CF UABC cultures showed CFTR_inh_-172-sensitive short-circuit currents of 13 ± 3 μA/cm^2^ (31 ± 5 % relative to non-CF) which was significantly higher than currents of 0.78 ± 0.04 μA/cm^2^ seen in uncorrected CF controls (p < 0.0001, t-test). Corrected ΔF508 compound heterozygous samples showed CFTR_inh_-172-sensitive short-circuit currents of 9 ± 2 μA/cm^2^ (20 ± 5 % relative to non-CF) which was significantly higher than uncorrected CF controls (p < 0.0001, t-test). Non-CF UABC cultures showed CFTR_inh_-172-sensitive short-circuit currents of 42 ± 6 μA/cm^2^.

Corrected ΔF508 homozygous HBECs showed CFTR_inh_-172-sensitive Cl^-^ currents of 10 ± 1 μA/cm^2^ (51 ± 3 % relative to non-CF) compared to 2 ± 1 μA/cm^2^ seen in uncorrected ΔF508 homozygous HBECs (p < 0.0001, t-test) and 18 ± 3 μA/cm^2^ seen in non-CF HBECs. Genotype information, percent alleles corrected and restoration of CFTR_inh_-172 short-circuit currents relative to average non-CF currents for individual samples are presented in Table 1.

### Gene Edited Basal Cells Can be Embedded in FDA-approved Porcine SIS membrane

As a path to transplanting *ex vivo* genetically corrected autologous cells into patients, we explored whether we could embed genetically edited airway cells on a pSIS membrane that is already in clinical use and FDA approved for several indications, including sinonasal repair (Nayak et al., 2018). We optimized the seeding density and culture conditions to obtain a monolayer of basal cells that retained expression of KRT5. We determined that the optimal plating density to achieve 50-70% primary cell coverage in four days was 100,000 cells/cm^2^ (Figure 3A). Hematoxylin and eosin (H&E) staining of pSIS in cross-section showed cells embedded as a monolayer (Figure 3B). Cells seeded on pSIS membrane remained KRT5+ (Figure 3C subject 1, Figure S5 A-I Subjects 2-4). Manders’ co-localization coefficients were calculated and the fraction of live (calcein green positive) cells also positive for KRT5 (M1) was determined to be 53 ± 15% (n = 4 from 4 individual donors). The fraction of KRT5+ cells corresponded to the KRT5+ fraction measured on the day of seeding did not change appreciably after proliferating on the pSIS membrane (Figure S5J). Thus, the pSIS membrane is a suitable scaffold to optimize transplantation of corrected cells in animal models.

**Figure 3:**
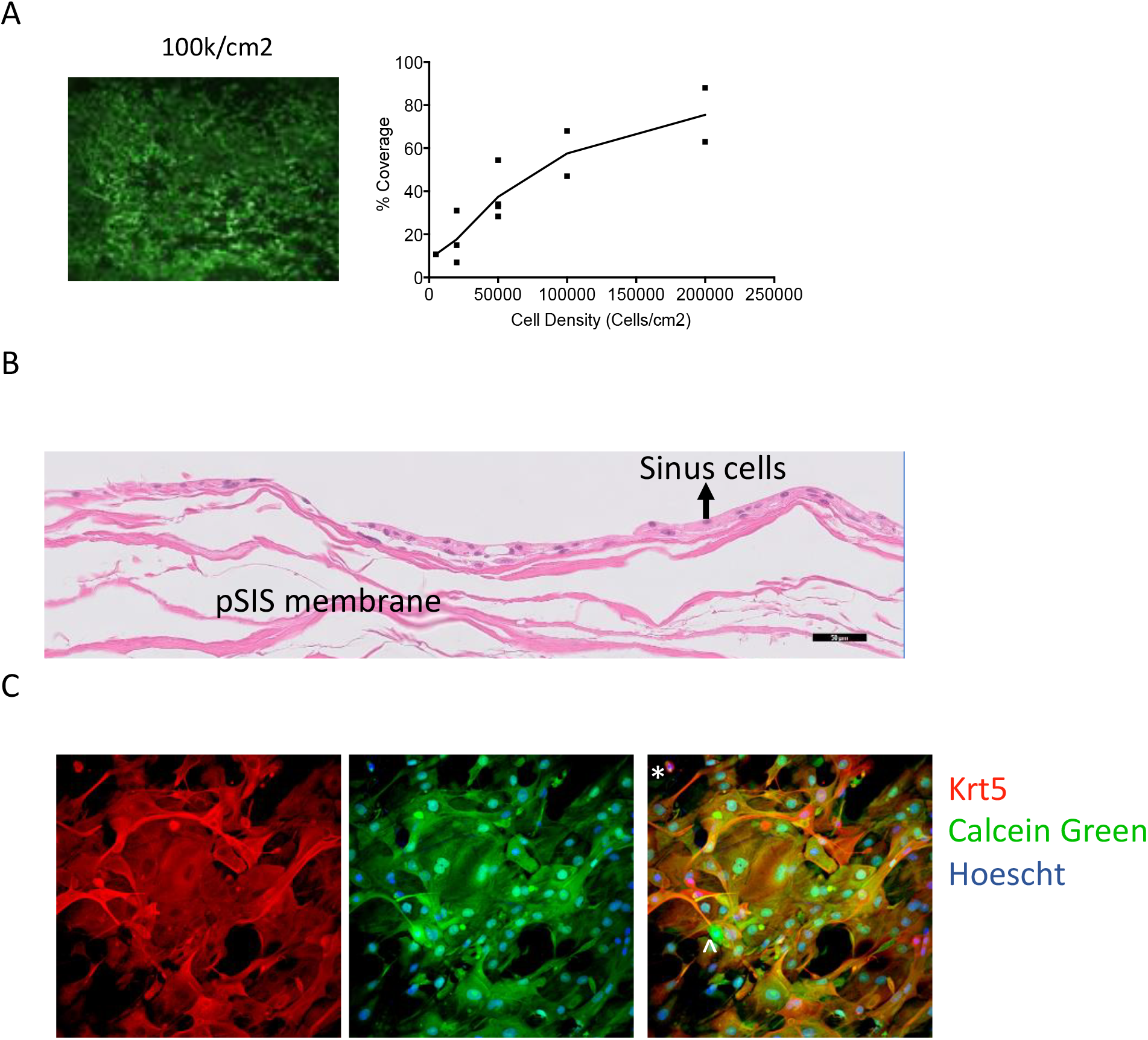
**(A)** Edited UABC cells plated on pSIS membranes at a density of 10^5^ cells/cm^2^ resulted in 50-70% confluence in four days **(B)** H&E staining shows a monolayer of cells on pSIS membranes (scale = 50 μm) **(C)** Sheets fixed on day 4 after embedding on pSIS membrane were KRT5+. Calcein green indicates live cells and KRT5+ cells are stained red. A few cells are positive for calcein green but not KRT5 (^). Some non-viable cells were still KRT5+ (*). Manders’ coefficients were calculated. The fraction of calcein green positive cells also positive for KRT5 was determined to be 78% for sample presented here (average = 53 ± 15% for n = 4 biological replicates).

## Discussion

The discovery of CF as a monogenic disease caused by CFTR mutations in 1989 prompted several attempts to use gene therapy for treatment (Griesenbach et al., 2015). These studies employed various viral and non-viral strategies but failed to show significant benefit (Griesenbach et al., 2015). The recent development of genome editing using Cas9 and other nucleases (e.g., zinc finger nucleases) has prompted a renewed effort to correct CF causing mutations. Although early studies achieved low levels of correction in intestinal stem cells and iPSCs (Crane et al., 2015; Firth et al., 2015; Schwank et al., 2013), correction of CFTR mutations at high efficiencies in therapeutically relevant primary airway stem cells has not previously been demonstrated.

In this study, we achieved correction of the ΔF508 mutation in ∼40% of alleles in primary upper airway and bronchial basal stem cells obtained from CF patients. This level of correction is a 100-fold improvement over previous studies using CRISPR/Cas9 and is within the range necessary for clinically significant restoration of CFTR function. Significantly, our approach achieves this high level of correction without the use of any drug-based selection strategy (e.g., puromycin), a key feature that is relevant for clinical translation (Schwank et al., 2013). When cultured in air-liquid interfaces, corrected UABC and HBECs gave rise to differentiated epithelia containing ciliated and mucus producing cells. Different media and culture conditions have been reported for the culture of epithelial sheets in air-liquid interface (Gentzsch et al., 2017). To ensure that we measured true CFTR function, we tested two commonly used media conditions (Pneumacult™ ALI medium and ALI medium reported previously (Gentzsch et al., 2017)). Although there were differences in the response to forskolin, the response to CFTR_inh_-172 was similar under both conditions in both non-CF and corrected CF cells (Figure S6). The data reported in Figure 2 are from cells cultured in Pneumacult™. The differences in the forskolin response is likely due to the presence of factors that increase baseline CFTR stimulation (e.g. cholera toxin) but this is unknown since the composition of Pneumacult™media is proprietary.

We observed restoration of CFTR function in both ΔF508 compound heterozygous as well as homozygous samples. However, since the sgRNA is located a few base pairs away from the ΔF508 mutation site and is active in non-CF cells, half of the replaced alleles did not contain the ΔF508 mutation. Therefore, as expected, a higher level of allelic correction (∼2-fold higher) is required in compound heterozygous samples for equivalent restoration of CFTR function. Previous studies have attempted to ascertain the minimal level of correction necessary to restore normal Cl^-^ transport by co-culturing non-CF or corrected CF cells and CF cells in ALI. These reports indicate that 10-50% normal cells are sufficient to restore non-CF level Cl^-^ transport in ΔF508 homozygotes (Johnson et al., 1992; Shah et al., 2016). Another study used a lentivirus based strategy to over-express CFTR under the control of an RSV promoter in pig epithelia (Cooney et al., 2016). They reported low genomic integration (<1 copy per 10 cells) but observed high levels of CFTR function. The minimal level of gene correction in the endogenous CFTR locus that can restore CFTR function to non-CF levels has not been previously reported. We detected as much as 2-fold difference in CFTR function in technical replicates from the same CF donor despite the same level of correction in cells from the same individuals (Table 1, Figure 2H). On average, 30-40% allelic correction restored CFTR function to 50% of non-CF levels in bronchial cells. However, one bronchial sample with 40% allelic correction showed 99% CFTR function relative to non-CF controls. Average CFTR function is approximately 60% higher in corrected HBECs (50% of non-CF levels) than UABCs (30% of non-CF levels). However, since the UABCs and HBECs were not from the same donors, it is unclear if the differences are due to underlying biology of these related but still distinct cells or due to differences in the genetic backgrounds of the individuals.

CFTR function has been reported to vary logarithmically in organ outputs measured *in vivo* (e.g. sweat chloride) and has been shown to be rate-limiting at low levels of CFTR expression (Char et al., 2014). Thus, even augmentation to a low level of CFTR function may provide significant clinical benefit. For example, patients homozygous for the R117H mutation experience infertility and mildly increased sweat chloride but are completely free of any respiratory or pancreatic symptoms (De Nooijer et al., 2011). Significantly, R117H and other class IV mutations are associated with significantly lower mortality compared to class II mutations such as ΔF508 (McKone et al., 2003). Patch clamp and apical conductance measurements on cells expressing exogenous R117H-CFTR showed as little as 15% Cl^-^ conductance relative to cells expressing wild-type CFTR (Sheppard et al., 1993). By way of contrast, Char et al. estimated <2% CFTR function relative to WT levels in patients with R117H mutations (Char et al., 2014). Thus, the 20-50% function relative to WT in CFTR function we found in our studies would be predicted to provide a meaningful clinical benefit to patients if achieved *in vivo*. This benefit may also be expected to be durable, suggested by our findings that targeting basal stem cells for gene correction does not impair their ability to contribute to differentiated airway epithelial cells.

Achieving correction of CFTR mutations *in vivo*, either by transplanting corrected cells or editing *in vivo*, remains a hurdle for gene therapy. Transplantation of airway stem cells into the lower airways have been reported in animal models but further optimization is necessary for clinical use (Rosen et al., 2015). We focused our experiments on upper airway basal cells since the ease of access to upper airway tissue may help clinical translation by providing readily accessible airway stem cells and by enabling surveillance of the sites after transplantation. Moreover, sinuses of CF patients have been shown to act as a reservoir for drug resistant pathogens that cause chronic lung infections (Hansen et al., 2012). Thus, chronic rhinosinusitis and recurrent infectious disease of the upper airways are unmet medical needs that affect CF patients. Chronic rhinosinusitis and recurrent infections are not ameliorated by lung transplantation and AAV mediated gene therapy in the maxillary sinus was unsuccessful (Wagner et al., 2002). The surface area of one maxillary sinus has been estimated to be ∼13 cm^2^ (Oliveria et al., 2014). Given that 10^5^ cells /cm^2^ on the pSIS membrane represents an optimal cell density, we estimate that 1.3 million corrected cells will be sufficient to completely cover the surface of one maxillary sinus even without any lateral outgrowth of gene-corrected basal cells following implantation. This cell yield is entirely achievable using our current approach, and transplantation with *in vivo* analysis will be the focus of our future studies.

## Conclusion

Our study shows that a platform consisting of Cas9, MS-sgRNA and AAV6 can be used to correct the ΔF508 mutation in primary airway stem/progenitor cells obtained from CF patients. This selection-free strategy achieves clinically significant restoration of CFTR function in differentiated epithelial sheets derived from corrected stem cells. The study paves the way for future experiments to optimize the transplantation of corrected UABCs into the upper airway to treat CF sinus disease. Successful optimization of stem cell transplantation into the sinuses may then enable further investigations on the use of cell therapies to treat CF lung disease.

## Experimental Methods

### EN media

ADMEM/F12 supplemented with B27 supplement, Nicotinamide (10 mM), human EGF (50 ng/mL), human Noggin (100 ng/mL), A83-01 (500 nM), N-acetylcysteine (1 mM) and HEPES (1 mM)

### Cell Culture

Upper airway tissue obtained from endoscopic surgery was cut into small pieces (1-2 mm^2^). Tissue pieces were washed with 10 ml sterile, PBS w/2X antibiotic/antimycotic (Penicillin, Streptomycin, Amphotericin B, Gibco # 15240062) on ice and digested with pronase (1.5 mg/mL, Sigma #P5147) for 2 h at 37 °C or at 4 °C overnight. Digestion was stopped using 10% FBS. Digested tissue was filtered through cell strainers (BD Falcon # 352350) into a sterile 50 ml conical tube. The mixture was centrifuged at 600xg for 3 minutes at room temperature. RBC lysis was then performed using RBC lysis buffer (eBioscience™) as per manufacturer’s instructions. After RBC lysis, cells were suspended in 1 mL EN media and counted. A small sample was fixed using 2% paraformaldehyde and permeabilized using Tris-buffered saline with 0.1% Tween 20. Cells were stained for cytokeratin 5 (KRT5, Abcam, ab 193895). An isotype control (Abcam, ab 199093) was used to control for non-specific staining. KRT5+ cells were plated at a density of 10,000 cells per cm^2^ as monolayers in tissue culture plates coated with 5% Matrigel. Cells were incubated at 37 °C in 5% O_2_ and 5% CO_2_ in EN media with 10 μM ROCK inhibitor (Y-27632, Santa Cruz, sc-281642A). Organoids were grown as 40 μl Matrigel domes submerged under identical medium at initial seeding density of KRT5+ 20,000 cells/dome. Cells obtained from CF patients were grown in EN media supplemented with additional antimicrobials for two days (Fluconazole – 2 μg/mL, Amphotericin B 1.25 μg/mL, Imipenem – 12.5 μg/mL, Ciprofloxacin – 40 μg/mL, Azithromycin – 50 μg/mL, Meropenem −50 μg/mL). The concentration of antimicrobials was decreased 50% after 2-3 days and then withdrawn after editing (day 5-6).

### Gene Editing

Cells were cultured in EN media with 10 μM ROCK inhibitor (Y-27632, Santa Cruz, sc-281642A). The presence of ROCK inhibitor for at least 24 h was critical for cell survival after electroporation. Media was replaced on day 3 and day 4 after plating from tissue. Gene correction was performed 5 days after plating. Cells were detached using TrypLE Express Enzyme (Gibco™12605010). Cells were resuspended in OPTI-MEM (Gibco™31985062) at a density of 5 million cells/mL. Other electroporation buffers were also tested but the best results were obtained with OPTI-MEM. A similar observation has been reported in intestinal organoids (Fujii et al., 2015). Electroporation (Nucleofection) was performed using Lonza 4D 16-well Nucleocuvette™ Strips (Lonza, V4XP-3032). 6 μg of Cas9 (Integrated DNA Technologies, IA, Cat: 1074182) and 3.2 μg of MS-sgRNA (Trilink Biotechnologies, CA) (molar ratio = 1:2.5) were complexed at room temperature for 10 minutes. 100,000 cells (20 μL of OPTI-MEM with 5 million cells/mL) was added to Cas9/MS-sgRNA mixture used per well and transferred to the strip. Cells were electroporated using the program CA-137. 80 μL of OPTI-MEM was added to each well after electroporation. Cells were transferred to a 12 well plate coated with 5% Matrigel (density = 20,000 cells/cm^2^) and 400 μL of EN media with 10 μM ROCK inhibitor was added. AAV carrying the correction template was added immediately after electroporation to maximize transduction (Bak et al., 2018; Charlesworth et al., 2018). Multiplicity of Infection (MOI) of 10^6^ particles per cell (as determined by qPCR) was optimal. AAV titers can also be determined by droplet digital PCR (ddPCR) but the output results in titers are 10-fold lower for the same sample when compared to qPCR. If AAV titer is measured using ddPCR, the appropriate MOI would be 10^5^ particles per cell. Media was replaced 48 h after electroporation. Gene correction levels were measured at least 4 days after electroporation.

### Measuring Gene Correction

4 days (or more) after gene correction, genomic DNA was extracted from cells using Quick Extract (Lucigen, QE09050) as per manufacturer’s instructions. The ΔF508 locus was amplified using the primers:

Forward: CCTTCTACTCAGTTTTAGTC

Reverse: TGGGTAGTGTGAAGGGTTCAT

The PCR product was sanger sequenced (primer: AGGCAAGTGAATCCTGAGCG) and the percent of corrected alleles was determined using TIDER.

### Measuring off-target activity

Primary UABCs from non-CF patients were electroporated with Cas9 and sgRNA without the HR template. gDNA was extracted 4-5 days after RNP delivery. Potential off-target sites were identified using the bioinformatic tool COSMID (Cradick et al., 2014) allowing for 3 mismatches within the 19 PAM proximal bases. Predicted off-target loci were initially enriched by locus specific PCR followed by a second round of PCR to introduce adaptor and index sequences for the Illumina MiSeq platform. All amplicons were normalized, pooled and quantified using a Qubit (ThermoFisher Scientific) and were sequenced using a MiSeq Illumina using 2 x 250bp paired end reads. INDELs at potential off-target sites were quantified as previously described (Lee et al., 2016).

### Air-Liquid Interface Culture of Corrected UABCs and HBECs

Gene corrected cells were plated 4-10 days after editing. 30,000 to 60,000 cells per well were plated in 6.5 mm Transwell plates with 0.4 μm pore polyester membrane insert. EN media was used to expand cells for 1-2 weeks. Once cells were confluent in Transwell inserts and stopped translocating fluid, media in the bottom compartment was replaced with Pneumacult ALI media. For comparison, a small batch of cells were also cultured in media obtained from the University of North Carolina (UNC media)(Gentzsch et al., 2017). The need for an additional coating of plates with collagen IV was also tested.

### Immunoblot

Immunoblotting methods were used to compare CFTR protein expression pre/post-correction. Calu-3 cells, non-CF nasal cells, and patient-derived ΔF508 homozygous pre/post-correction were plated and cultured according to the above methods. Lysis was performed by incubating cells for 15 minutes in ice-cold RIPA buffer supplemented with EDTA-free protease inhibitor. Lysates were gathered and rotated at 4°C for 30 min, then spun at 10,000 × g at 4°C for 10 minutes to pellet insoluble genomic material. The supernatant was collected and mixed with 2x Laemmli sample buffer containing 100 mM DTT and subsequently heated at 37°C for 30 min. Approximately 5 μg of total protein was loaded for Calu-3 lane, and approximately 14 μg of total protein was loaded for non-CF nasal and ΔF508 cell lines and fractionated by SDS-PAGE, then transferred onto PVDF membrane. Blocking was performed with 5% nonfat milk in TBST (10 mM Tris, pH 8.0, 150 mM NaCl, 0.5% Tween 20) for 60 min. The membrane was probed with antibodies against CFTR (Ab450, 1:1000), and β-actin (1:10,000). Membranes were washed and incubated with a 1:10,000 dilution of horseradish peroxidase-conjugated anti-mouse for 1 h, then developed by SuperSignal™ West Femto Maximum Sensitivity Substrate (Thermo Fisher Scientific, 34095).

### Ussing Chamber Functional Assays

Ussing chamber measurements were performed 3-5 weeks after cells had stopped translocating fluid as described before. For Cl^-^ secretion experiments with UABCs, and HBECs, solutions were as following in mM: Mucosal: NaGluconate 120, NaHCO_3_ 25, KH_2_PO_4_ 3.3, K_2_HPO_4_ 0.8, Ca(Gluconate)^2^ 4, Mg(Gluconate)^2^ 1.2, Mannitol 10; Serosal: NaCl 120, NaHCO_3_, 25, KH_2_PO_4_ 3.3, K_2_HPO_4_ 0.8, CaCl_2_ 1.2, MgCl_2_ 1.2, Glucose 10. The concentration of ion channel activators and inhibitors were as follows:

Amiloride -10 μM – Mucosal

Forskolin -10 μM – Bilateral

VX-770 -10 μM – Mucosal

CFTR_inh_-172 -20 μM – Mucosal

UTP – 100 μM – Mucosal

### Embedding cells on pSIS membrane

pSIS membranes (Biodesign^®^ Sinonasal Repair Graft; COOK Medical, Bloomington, IN) were placed in 8 well confocal chambers. UABCs were seeded 4-8 days after electroporation. Four days after seeding, cells were incubated with calcein green for 15 min and the excess washed away. Cells were imaged using a dissection scope to identify densities that provided optimal coverage. pSIS membranes with cells were fixed with 4% paraformaldehyde, permeabilized with TBS-T (0.1% Tween 20) and stained for KRT5 (ab193895) and imaged using Leica SP8 confocal microscope.

### Key resources

**Key resources:**
| Reagent or Resource | Source | Identifier |
| --- | --- | --- |
| **Antibodies** |  |  |
| Anti-Cytokeratin 5 antibody [EP1601Y] (Alexa Fluor® 647) | Abcam | ab193895 |
| Rabbit IgG, monoclonal [EPR25A] - Isotype Control (Alexa Fluor® 647) | Abcam | ab199093 |
| Alexa Fluor® 488 anti-human/mouse CD49f Antibody | Biolegend | 313608 |
| **Biological Samples** |  |  |
| Upper airway basal cells (UABCs) | Stanford University<br>University of Arizona | N/A |
| Bronchial epithelial cells (HBECs) | Lonza, McGill University | N/A |
| **Reagents** |  |  |
| Adeno-associated Virus Serotype 6 with ΔF508 correction template | Vigene Inc | Custom order |
| Adeno-associated Virus Serotypes 1,2,5,6,7,8,9, DJ with UBC-GFP | Vigene Inc | Custom Order |
| Alt-R® S.p. Cas9 Nuclease 3NLS, 500 µg | Integrated DNA technologies | 1074182 |
| Primers | Integrated DNA technologies | N/A |
| MS-sgRNA | Trilink Biotechnologies | O-5200 |
| Y-27632 dihydrochloride | Santacruz Biotechnology | sc-281642A |
| HEPES | Gibco | 15630-080 |
| GlutaMAX-1 (100x) | Gibco | 35050-061 |
| NIC (Nicotinamide) | Sigma | N0636 |
| B-27 supplement (50x) | Gibco | 125870-01 |
| A83-01 | Torcis | 2939 |
| Human EGF | R&D | 236-EG-01M |
| Human Noggin | R&D | 6057-NG/CF |
| admem F-12 | Invitrogen | 12634-028 |
| Biodesign™ Sinonasal Graft | Cook Medical | G35950 |
| CellTrace™ Calcein Green, AM | Invitrogen | C34852 |
| Anti-CFTR antibody 450 | University of North Carolina | N/A |
| Pneumacult™ ALI media | Stem Cell Technologies | #05001 |
| Pneumacult™ Ex-Plus | Stem Cell Technologies | 05040 |

## Acknowledgements

We thank Dr. Scott Randell (University of North Carolina) for guidance on Ussing chamber assays and training on air-liquid interface cultures. We thank Dr. Jeffrey Beekman and Dr. Gimano Amatngalim for their guidance on Ussing chamber assays. The work was funded by grants from the California Institute of Regenerative Medicine (DISC2-09637), Stanford-SPARK MCHRI and the Cancer Prevention and Research Institute of Texas (RR14008 and RP170721).

---

**Figure S1.**
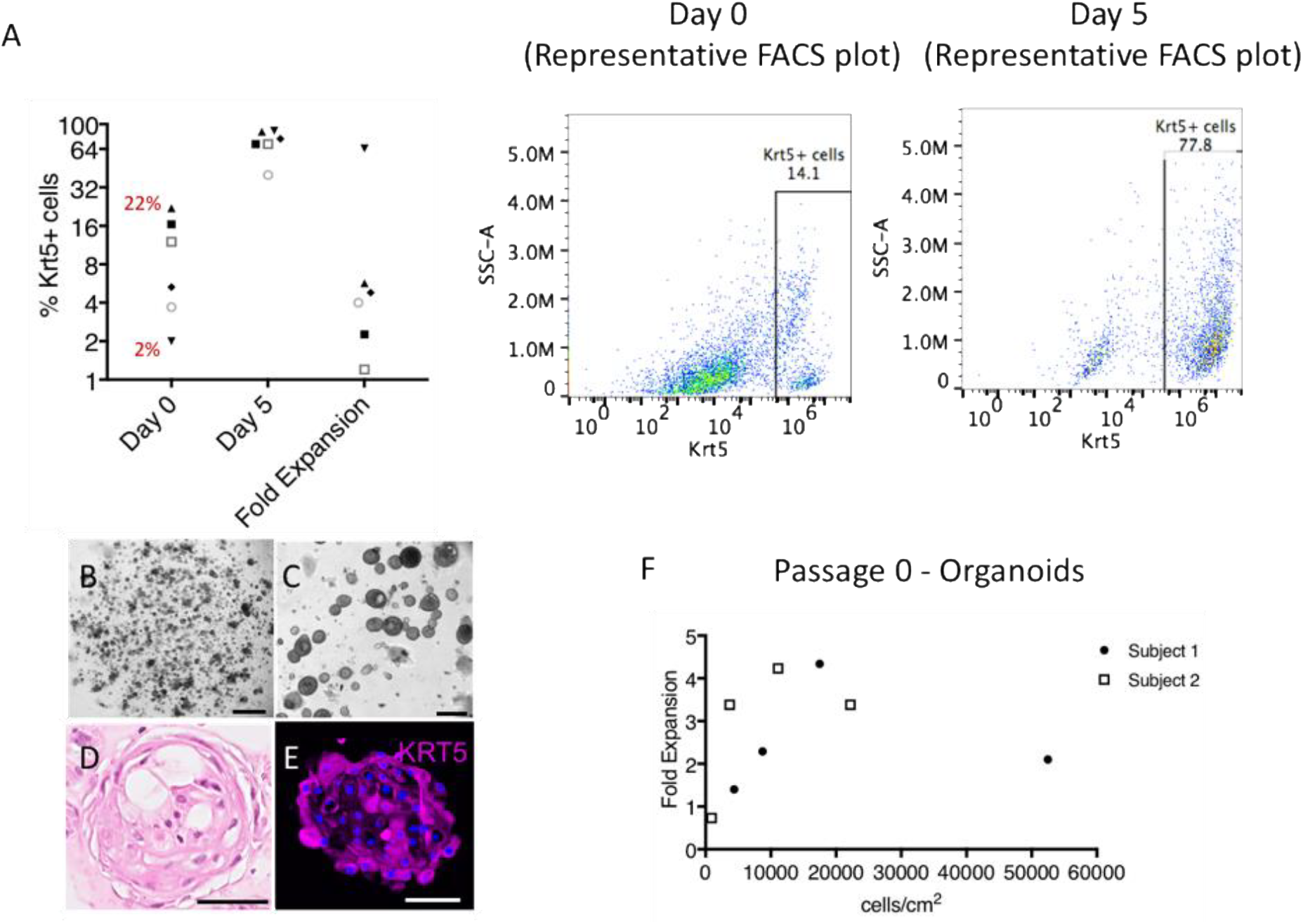
Characterization of human UABC organoids. **(A)** Percent KRT5+ cells on day 0 (2-22%, average ± SD = 10 ± 8%), day 5 (40-90%, average ± SD = 72 ± 18%) and fold expansion observed in 6 subjects (2-64%). Representative FACS plots of UABC organoids with KRT5+ gating on day 0 versus day 5. **(B-E)** Primary human UABCs were cultured as organoids in Matrigel domes in EN medium (A, scale = 1 mm and B, scale = 0.2 mm). Organoids were assessed by H&E staining (C, scale = 50 μm). Organoids were positive for KRT5 immunofluorescence (D, day 5, scale = 50 μm) at culture day 5. **(F)** Optimal organoid proliferation was observed at cells densities between 10,000-20,000 cells/cm^2^ at passage 0.

**Figure S2.**
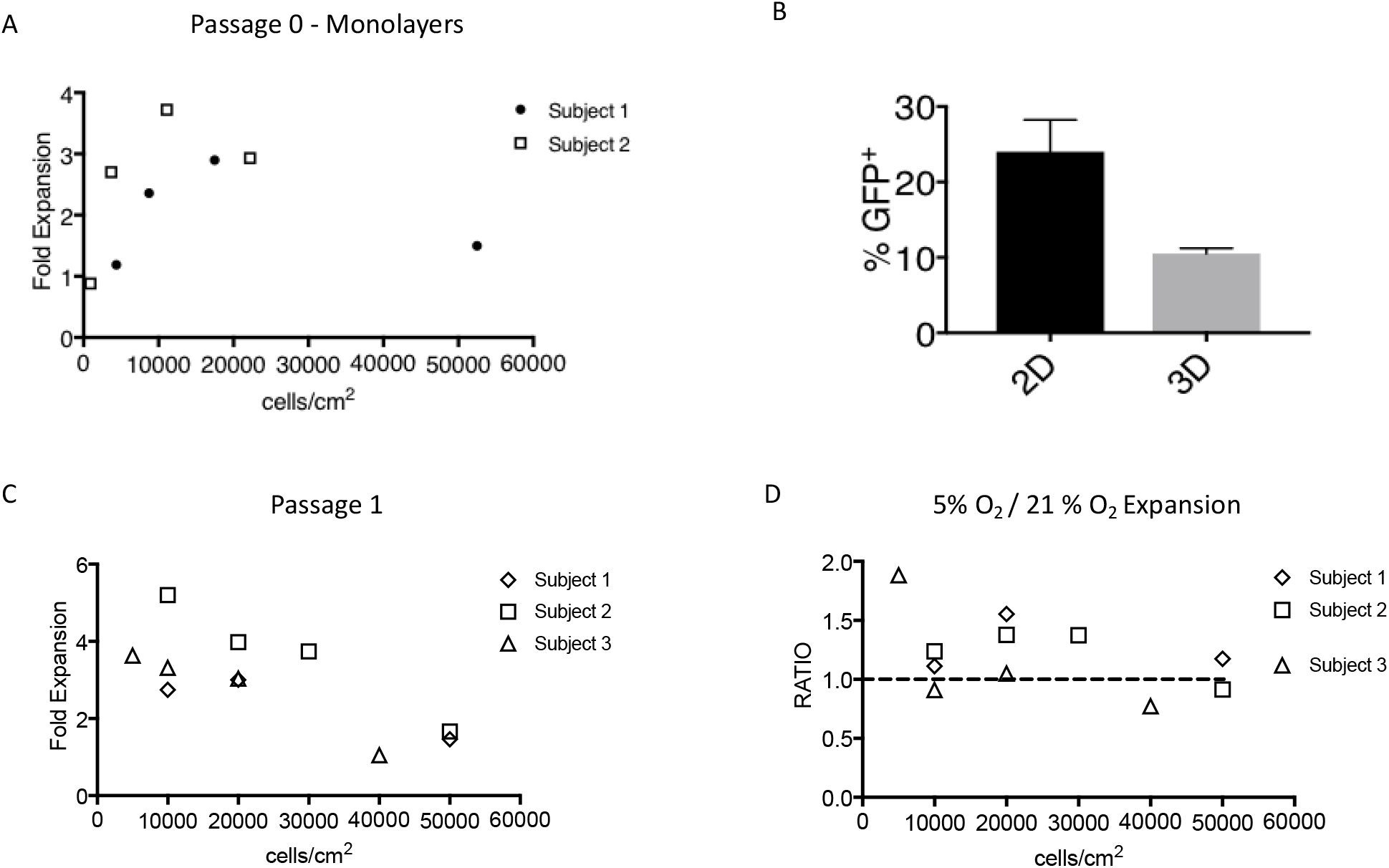
Improved gene editing in UABC monolayers versus 3D organoids. **(A)** Primary human UABCs were cultured as monolayers in tissue culture treated plates coated with 5% Matrigel in EN medium. Optimal organoid proliferation was observed at cells densities between 10,000-20,000 cells/cm^2^ at passage 0. **(B)** Cells cultured both as monolayers and organoids were edited to express SFFV/GFP at the CCR5 locus using a previously reported construct (Hendel et al., 2015). Editing efficiencies were higher for cells cultured as monolayers (N = 3, p = 0.02 by paired t-test) **(C)** Optimal cell density was also assessed in passaged cells and cell densities between 10,000-20,000 cells/cm^2^ were optimal at passage 1. **(D)** Culturing UABC monolayers in 5% O_2_ improved proliferation of cells from 2 out of 3 subjects (subjects 1 and 2).

**Figure S3:**
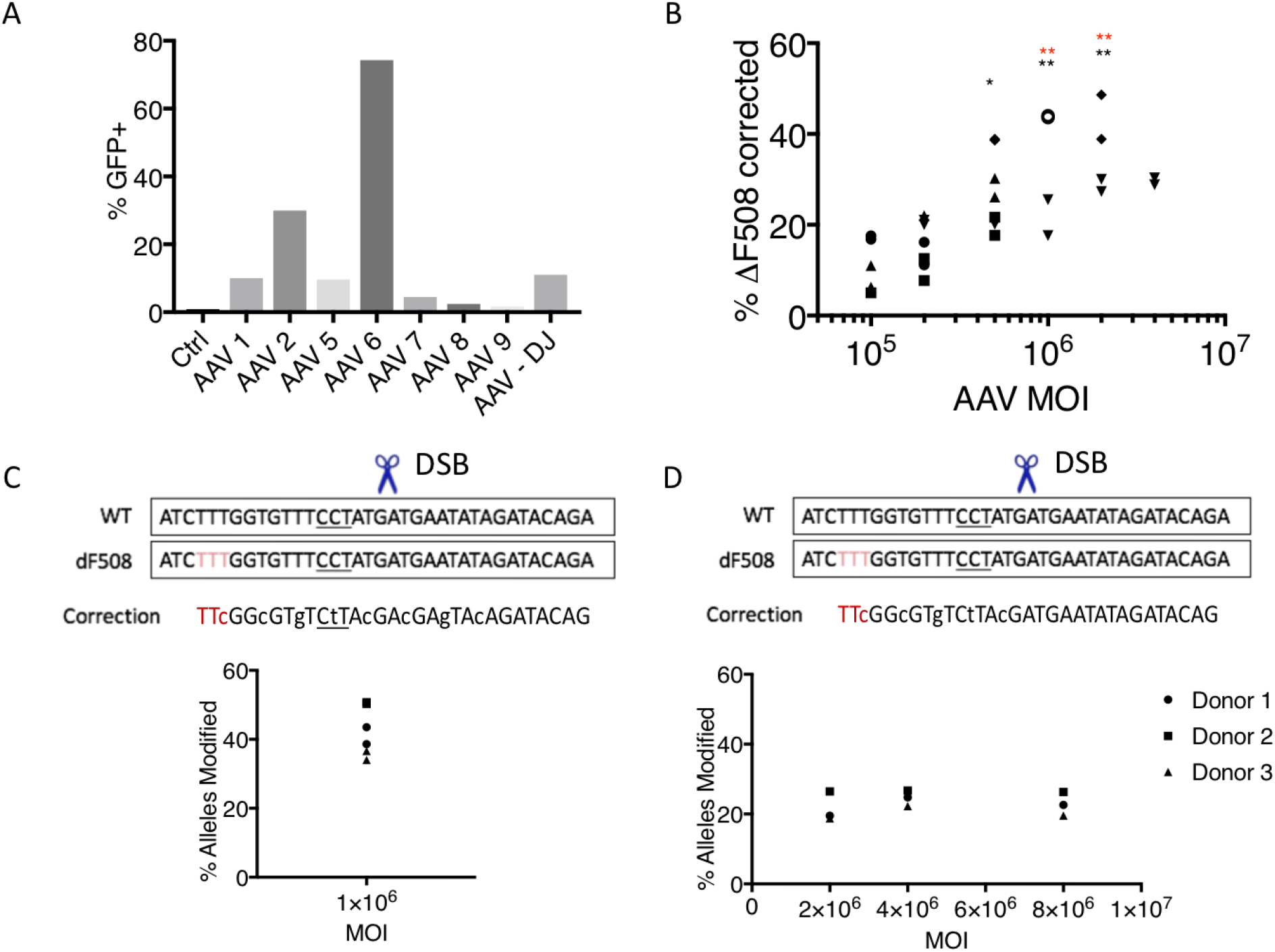
**(A)** AAV6 showed the best transduction in upper airway basal cells (UABCs). UABCs were transduced within 5 min after electroporation **(B)** In UABCs obtained from non-CF patients, MOIs of 10^6^ and 2×10^6^M vector genomes (vg) /cell showed significantly higher editing compared to MOIs < 2×10^5^ vg/cell. Different symbols represent cells from a different donor. **(C-D)** Two different correction templates with varying amounts of silent mutations (lower case letters) around the DSB site (scissors) were tested. PAM sequence is underlined. Templates with silent mutations on both sides of the DSB site (C) resulted in higher HR than templates containing mutations on one side (D).

**Figure S4:**
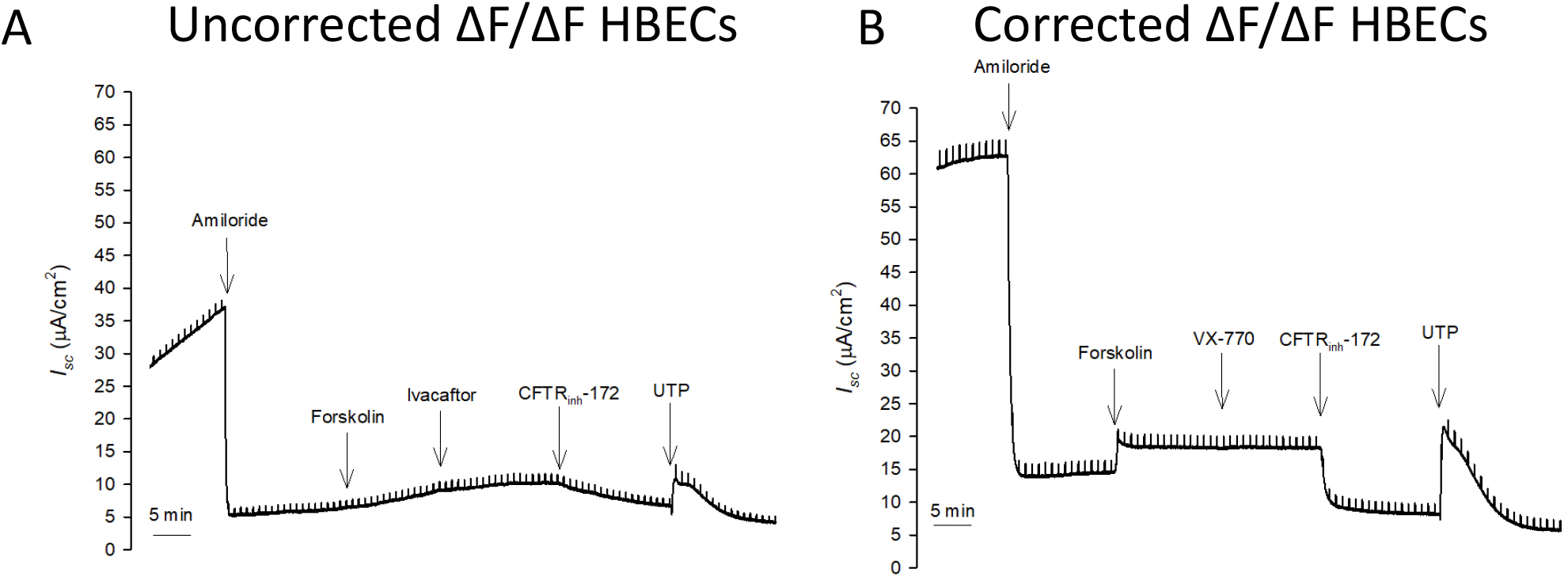
**(A)** Representative traces obtained from epithelial monolayers derived from ΔF508 homozygous (ΔF/ΔF) CF HBECs; **(B)** Correction of ΔF508 mutation in 30% of alleles resulted in a restoration of CFTR function

**Figure S5:**
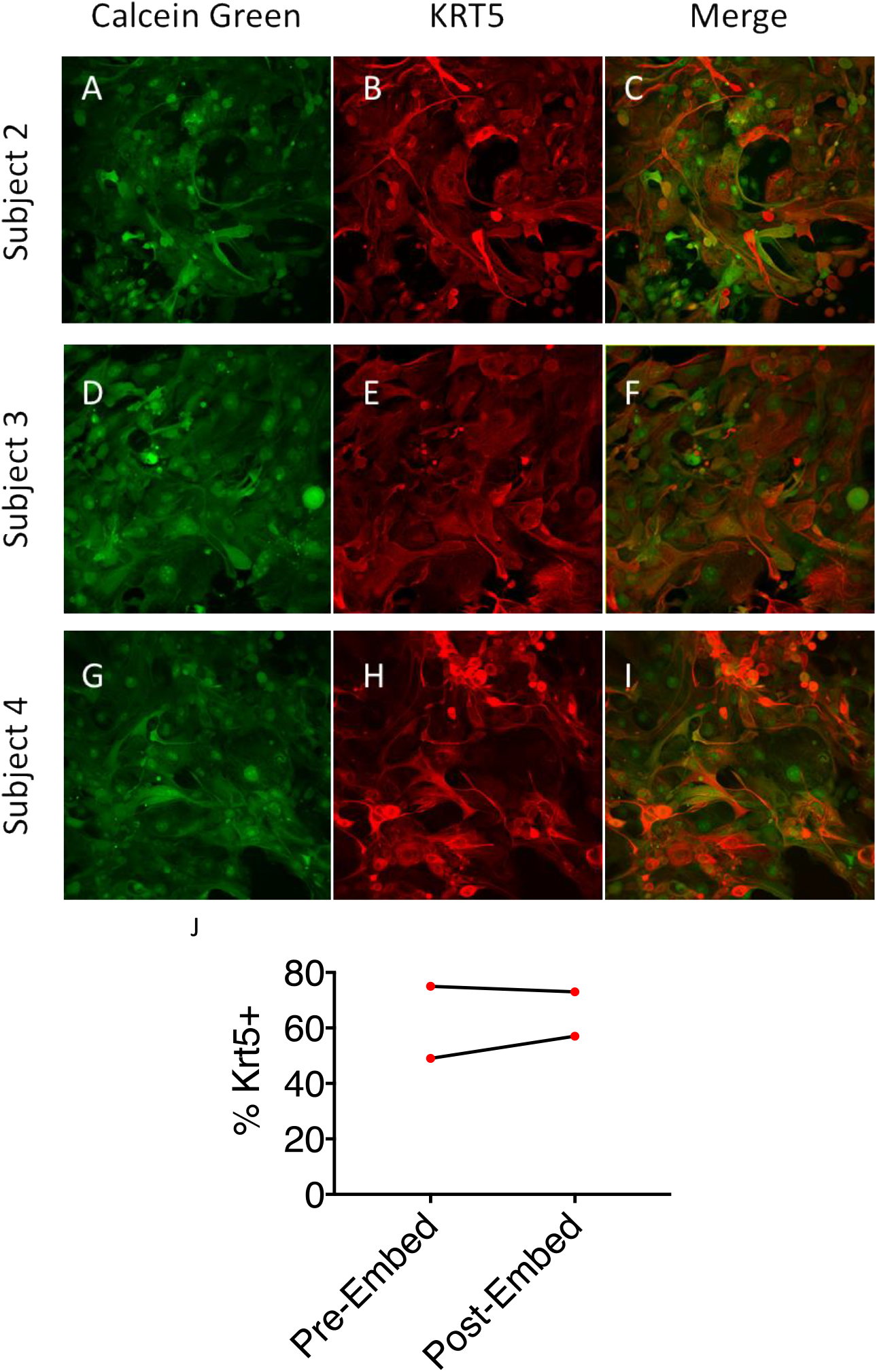
**(A-I)** Sheets fixed on day 4 after embedding on pSIS membrane were KRT5+. Calcein green indicates live cells and KRT5+ cells are stained red. Images from 3 additional subjects are shown here. (**J)** The fraction of KRT5+ cells was measured by flow cytometry on the day of embedding in pSIS membranes (Pre-embed). The fraction of calcein green positive cells that were also KRT5+ on day 4 after embedding was determined using confocal microscopy (Post-embed). The fraction of KRT5+ cells were similar indicating that edited cells did not experience a proliferative disadvantage. Thus, the pSIS membrane is a suitable scaffold to optimize transplantation of corrected cells in animal models.

**Figure S6:**
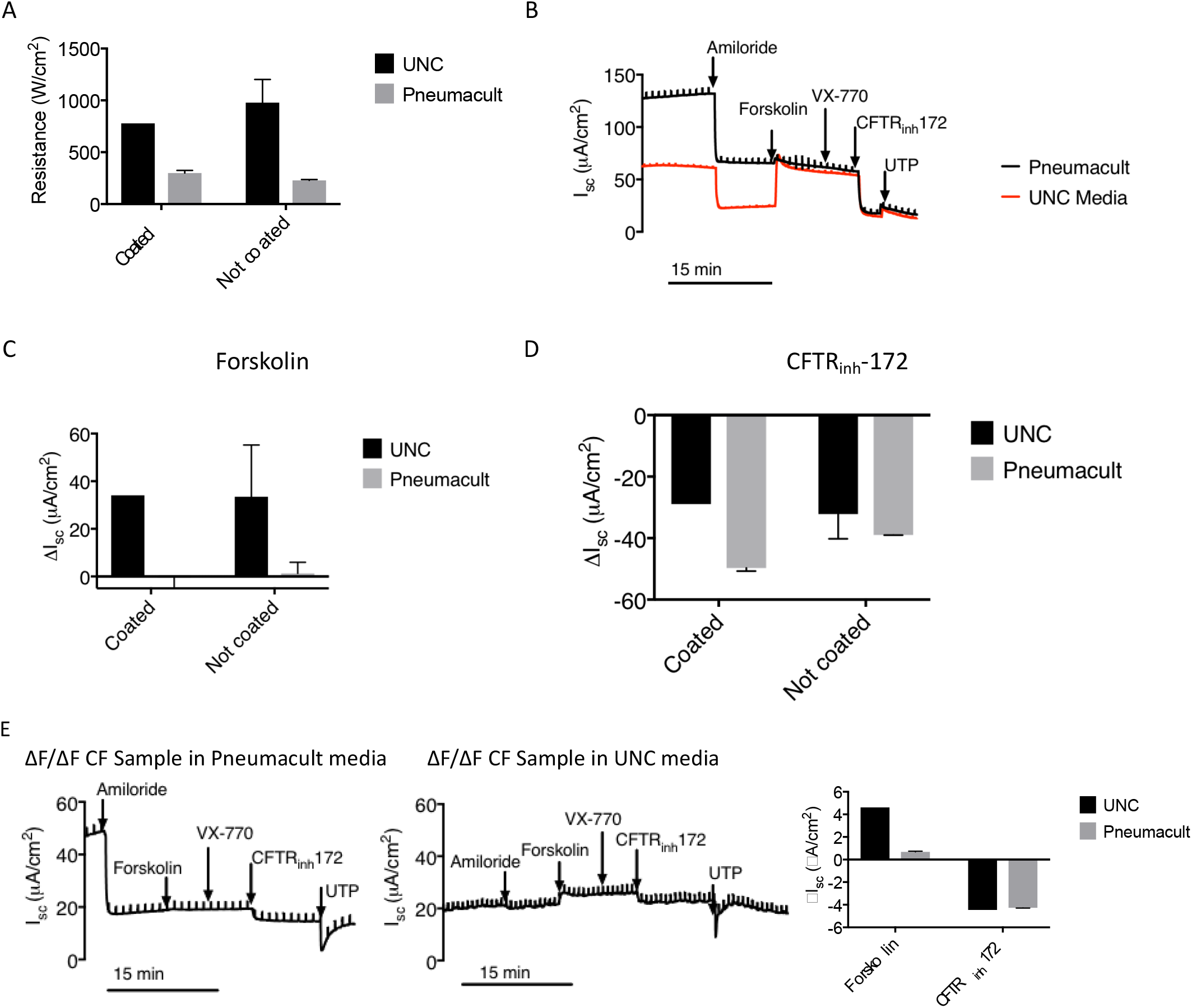
**(A)** UABCs from non-CF patients cultured in UNC media showed higher transepithelial resistances after differentiation on ALI **(B)** Representative traces from epithelial monolayers cultured in Pneumacult™ ALI and UNC media. Monolayers cultured in UNC media showed a more pronounced forskolin response. **(C)** In non-CF samples, short-circuit currents in response to forskolin were higher in monolayers cultured in UNC media. The presence or absence of collagen IV coating did not affect forskolin responses **(D)** Responses to CFTR_inh_-172 were similar between monolayers cultured in UNC media and Pneumacult ALI™ for non-CF cells. The presence or absence of collagen IV coating did not affect CFTR_inh_-172 responses **(E)** Upper airway basal cells from ΔF508 homozygous patient were edited (27% allelic correction) and differentiated using Pneumacult ALI™ and UNC media (no collagen IV coating) on ALI membranes. Similar to non-CF UABCs (C-D), responses to CFTR_inh_-172 were similar between monolayers cultured in UNC media and Pneumacult™ ALI but monolayers cultured in UNC media showed higher forskolin-stimulated short-circuit currents.

**Table S1:**
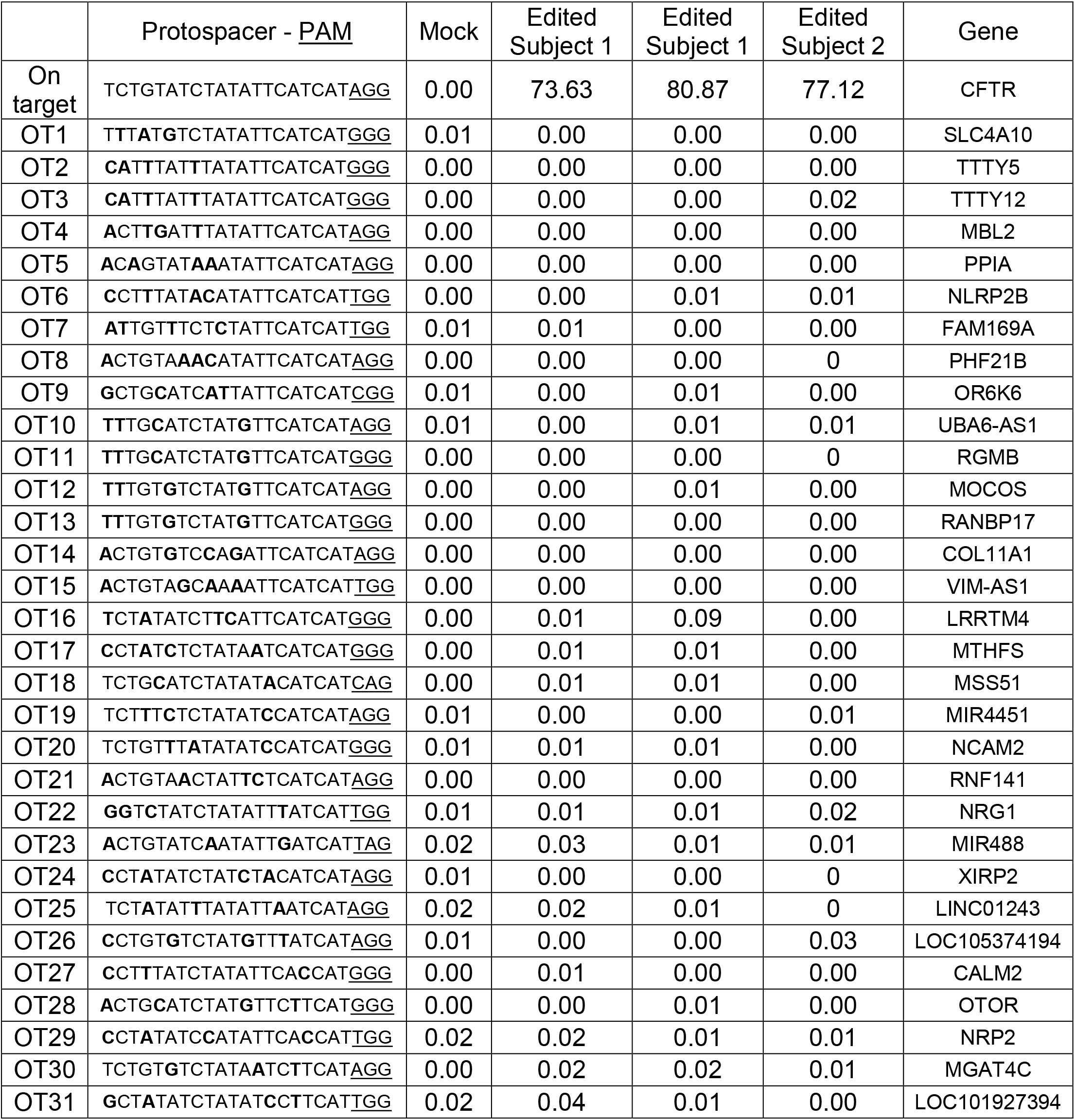
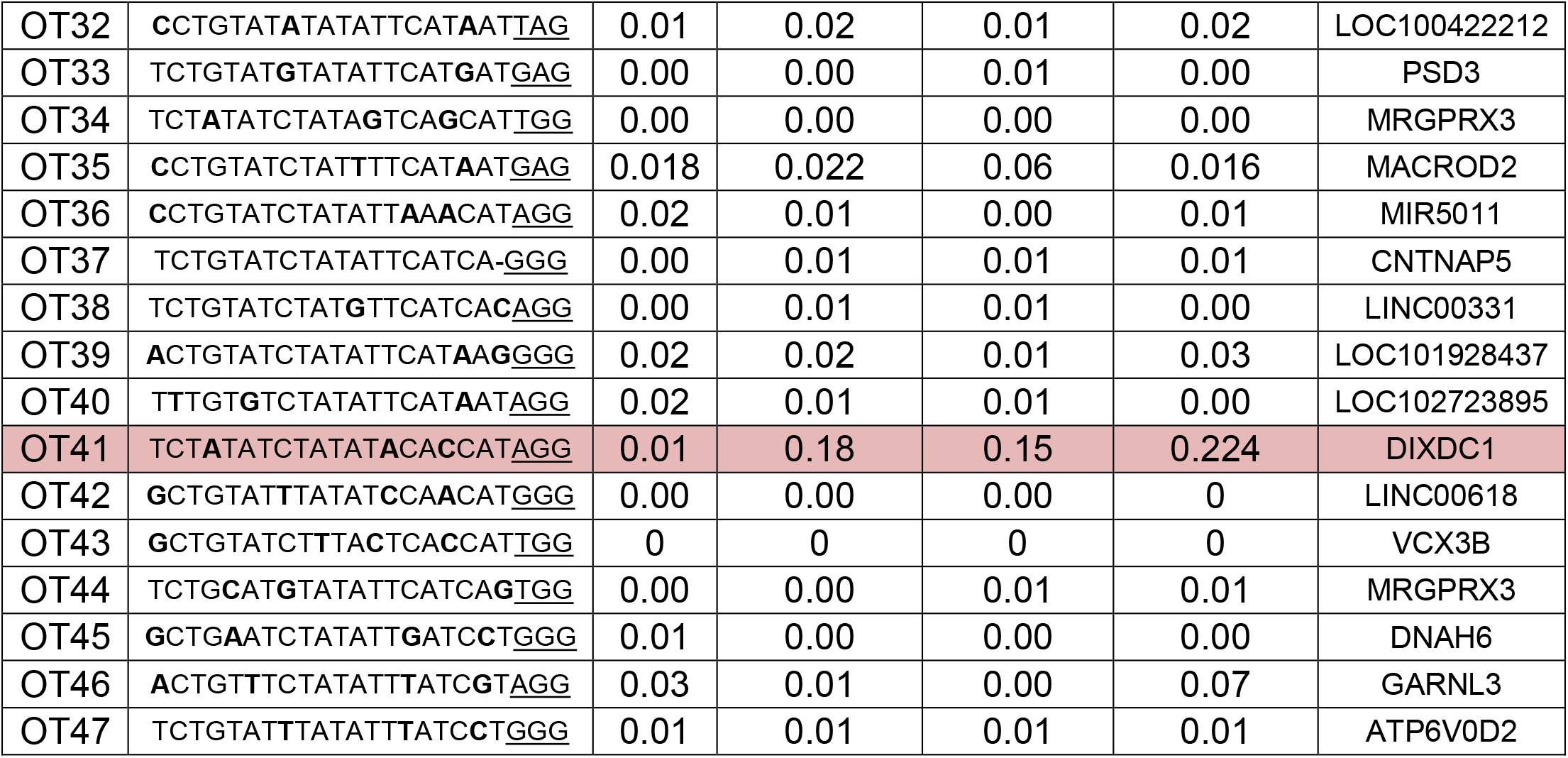
Off-target Activity of MS-SgRNA. Cas9 can induce double stranded breaks even if there are a few mismatches between the target genomic DNA and sgRNA. As a result, undesired insertions and deletions at off-target (OT) sites can happen. The MS-sgRNA targeting the ΔF508 locus shows very low levels of OT activity (< 0.2%) at one locus (OT-41) and high levels of on target activity. Mismatches are highlighted in bold and the protospacer adjacent motif (PAM) is underlined.

